# Artificially Inducing Close Apposition of Endoplasmic Reticulum and Mitochondria Induces Mitochondrial Fragmentation

**DOI:** 10.1101/005645

**Authors:** Victoria J. Miller, David J. Stephens

## Abstract

Cycles of mitochondrial fission and fission are essential for normal cell physiology. Defects in the machinery controlling these processes lead to neurodegenerative disease. While we are beginning to understand the machinery that drives fission, our knowledge of the spatial and temporal control of this event is lacking. Here we use a rapamycin-inducible heterodimerization system comprising both ER and mitochondrial transmembrane components to bring the ER membrane into close physical proximity with mitochondria. We show that this artificial apposition of membranes is sufficient to cause rapid mitochondrial fragmentation. Resulting mitochondrial fragments are shown to be distinct entities using fluorescence recovery after photobleaching. We also show that these fragments retain a mitochondrial membrane potential. In contrast, inducible tethering of the peripheral ER exit site protein TFG does not cause mitochondrial fragmentation suggesting that very close apposition of the two membranes is required.

## Introduction

Mitochondria routinely under-go cycles of fission and fusion (Bereiter-Hahn and Voth, 1994; Hermann and Shaw, 1998). Although a traditional depiction of a cell shows individual isolated mitochondria, in live cells they form a dynamic and connected tubular network that extends through-out the cytoplasm. Mitochondria provide energy for cells through the generation of ATP and have important roles in other cellular functions including apoptosis and autophagy (Nunnari and Suomalainen, 2012). Regulation of mitochondrial fission and fusion is important both for the normal physiology of the cell and for quality control of mitochondrial, with damaged mitochondria being removed by mitophagy (Yang et al., 2008; Youle and van der Bliek, 2012). Altering the balance of mitochondrial fission and fusion is sufficient to trigger cell cycle defects (Qian et al., 2012) and mutations in several of the genes encoding the machinery involved are linked to neurodegenerative diseases (Alexander et al., 2000; Delettre et al., 2000; Deng et al., 2008; Waterham et al., 2007; Zuchner et al., 2004).

In yeast, contacts between the ER and mitochondria are facilitated by the ER-Mitochondria Encounter Structure (ERMES) (Kornmann et al., 2009). These sites are required for mitophagy in yeast (Bockler and Westermann, 2014). However, the role of this complex in some aspects of mitochondrial metabolism is unclear (Nguyen et al., 2012) and no such stable structure has been defined in mammalian cells. ER tubules mark sites of mitochondrial divisions in both yeast and higher eukaryotes (Friedman et al., 2011). From this and other work, it has been proposed that wrapping of the ER membrane around the mitochondrion may generate initial constriction prior to the recruitment and action of the fission machinery including dynamin-related protein Drp1 (Lackner and Nunnari, 2009). More recent work has identified roles for actin through the action of the ER-localized formin INF2 (Korobova et al., 2013) and for myosin-2 (Korobova et al., 2014) in mitochondrial fission. A role has also been proposed for ER-mitochondrial contacts in the initiation of autophagy (Korobova et al., 2013) and more specifically mitophagy (Zuchner et al., 2004). COPII-coated secretory cargo exit sites on the ER membrane (ERES (Brandizzi and Barlowe, 2013)) have been linked to the initiation of autophagy (Tan et al., 2013) as have the membranes of the ERGIC which lie in close apposition to ERES (Ge et al., 2013). Consequently, one can imagine a scenario in which ERES spatially organize the apposition of ER and mitochondrial membranes and coordinate this with the initiation of mitophagy. TFG is a key component of the ERES and importantly appears to act as a mesh around newly formed COPII vesicles to maintain a structural integrity to these sites (Witte et al., 2011).

Given the extensive spread of both ER and mitochondria throughout most cells, the tight control of the role of ER-mitochondrial contact in driving fission is essential. How contact between these organelles drives fission versus other known functions such as calcium homeostasis or lipid transfer is also unclear. It is also unclear how some contact sites might exist to facilitate specific functions while others trigger mitochondrial fission. Indeed, the mechanisms that control how the number and location of sites of mitochondrial fission are dictated are not known.

Rapamycin-inducible heterodimerization uses rapamycin-binding domains to ectopically join two proteins together. It is most commonly used to sequester proteins away from their normal site of action (Inoue et al., 2005; Robinson et al., 2010). Here we have applied the ‘knock-sideways’ system to test whether driving the ER membrane into close proximity to the mitochondrial membrane is sufficient to direct mitochondrial fisison. We observe a rapid fission of mitochondria (initial constriction of mitochondria is observed within 60 seconds progressing to fragmentation from 20 minutes after the addition of rapamycin) consistent with a direct role for the close apposition of these membranes in triggering this event. This inducible system provides a means to interrogate the mechanisms that control the location and activity mitochondrial fission machinery in mammalian cells and to examine the wider function of ER-mitochondrial contacts in the cell.

## Results

We anchored the rapamycin-binding domain of FKBP12 to the cytoplasmic face of the ER membrane using a single transmembrane domain of 17 uncharged amino acids (Fig. 1A) (Bulbarelli et al., 2002; Ronchi et al., 2008). When expressed in cells this construct, which we have called FKBP-ER17, localizes to the ER membrane with a distribution indistinguishable from that of the related GFP-FP17 (FP for fluorescent protein (Ronchi et al., 2008)) (Fig. 1B). Rapamycin-dependent dimerization of the FKBP domain with the FKBP and rapamycin-binding (FRB) domain from mTOR is used in the knocksideways system to retarget FKBP-fusion proteins to the mitochondria. This is achieved through expression of a mito-YFP-FRB fusion that is constitutively associated with mitochondria by virtue of the import signal of outer mitochondrial membrane protein Tom70 (Robinson et al., 2010). Incorporation of YFP into this fusion allows visualization of mitochondria and the fate of the FRB fusion during these experiments. Here (Fig. 1C), we have engineered the system to drive close apposition of ER membranes (through FKBP-FP17) with mitochondrial membranes (with mito-YFP-FRB).

**Figure 1.**
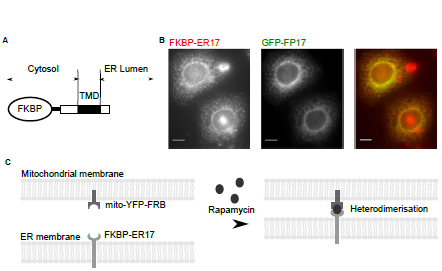
Schematic of engineered ER-mitochondrial contact. (A) Schematic of FKBP-ER17 construct, adapted from (Bulbarelli et al., 2002). The FKBP domain (oval) is linked to a transmembrane domain (TM) flanked by sequences up- and downstream. (B) Immunofluorescence labelling of methanol fixed HeLa cells co-expressing GFP-FP17 and FKBP-ER17. (C) Schematic of rapamycin-induced heterodimerization of mito-YFP-FRB and FKBP-ER17 bringing the ER and mitochondrial membranes together. Scale bar = 10 μm.

Cells were imaged following the addition of rapamycin to determine the effect of directing close apposition of ER and mitochondrial membranes on mitochondrial morphology. Fig 2A shows maximum intensity projections of cells stained with MitoTracker^®^ Red CMXRos (referred to hereafter as MitoTracker) immediately before and 49 minutes after incubation with 200 nM rapamycin. In some cases fragmented mitochondria formed a characteristic ‘doughnut’ shape. This is particularly evident in the enlargements (Fig 2A, insets). This change in mitochondrial morphology required that cells were transfected with plasmids expressing both mito-YFP-FRB and FKBP-ER17; Fig. 2B shows a non-expressing cell from the same field (with no visible mito-YFP-FRB protein) is shown as a control for the effects of both the rapamycin and the imaging procedure.

In the absence of rapamycin, mitochondria in both cells form a linked network (Fig. 2A and B showing a maximum intensity projection and Fig 2C showing a single z-plane). After the addition of rapamycin the mitochondria in the transfected cell fragment (initial constriction of mitochondria is observed within 60 seconds, with fragmentation apparent from 20 minutes after the addition of rapamycin), while the mitochondrial network in the control cell remains intact. Time-lapse images were taken of this process (Fig. 2D showing enlargements of the boxed region in Fig. 2C; see also Movie 1). These data show clear constriction of the mitochondria followed by apparent fission (Fig. 2D, arrow marks a mitochondrion post-fission).

**Figure 2.**
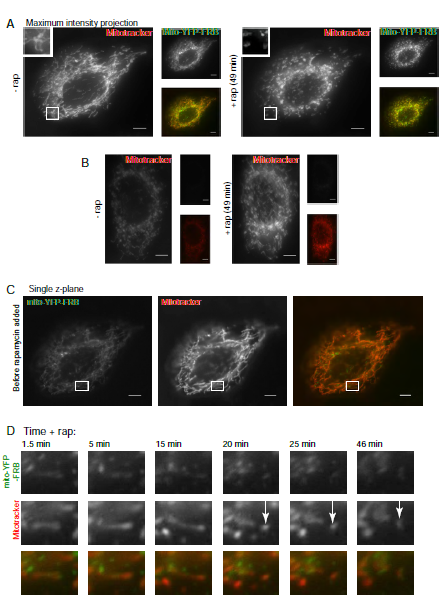
Mitochondrial fragmentation following inducible tethering of the ER membrane to mitochondria. (A) Maximal intensity projections of live cell imaging of HeLa cells co-transfected with FKBP-FP17 and mito-YFP-FRB before and after incubation with 200 nM rapamycin. Mitochondria are additionally stained with MitoTracker-Red. Images of both cells are taken from a single field of view, right-hand panel shows cell not expressing mito-YFP-FRB. (B) Single-plane frames from time-lapse imaging of cells shown in (A). Arrow indicates mitochondrial breakage. (C) Single-plane frames from time-lapse imaging of cells. Arrows show rapid clustering of mito-YFP-FRB on addition of rapamycin. Scale bars = 10 μm.

Immediately after addition of rapamycin, we observed that the mito-YFP-FRB, which is freely diffusible within the mitochondrial outer membrane, ‘clustered’ into discrete patches on the mitochondria surface within seconds of the addition of rapamycin (Fig. 3A and B and Movie 2). We conclude that bringing the ER and mitochondria into close proximity results in constriction of the mitochondrial membrane and subsequently results in mitochondrial fission.

**Figure 3.**
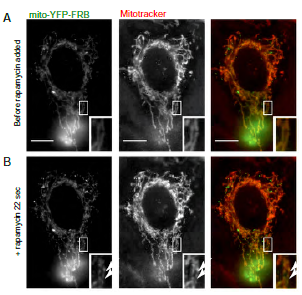
Addition of CCCP causes clustering of mito-YFP signal. HeLa cells co-transfected with FKBP-FP17 and mito-YFP-FRB before (A) and after (B) incubation with 200 nM rapamycin. Mitochondria are additionally stained with MitoTracker-Red. Single-plane frames from time-lapse imaging of cells prior to (A) and (B) immediately following addition of 200 mM rapamycin. Arrows show clustering of mito-YFP-FRB. Scale bar = 10 μm.

Damage leading to depolarization can also cause mitochondrial fragmentation (Benard et al., 2007; Hackenbrock, 1966; Hackenbrock, 1968). Fluorescence of the MitoTracker dye that we used here is dependent on the mitochondrial membrane potential. Live cell imaging confirmed that artificial mitochondrial fragmentation using this system was not associated with a loss of mitochondrial membrane potential. We added the uncoupler carbonyl cyanide ***m***-chlorophenylhydrazone (CCCP; Fig. 4, Movie 3) 49 minutes following the addition of rapamycin. The cell with fragmented mitochondria shows a rapid decrease in MitoTracker signal in the red channel (starting 4 minutes after the addition of 100 nM CCCP, with no further decrease in signal after 7.5 minutes) while the mito-YFP-FRP signal is unchanged. The control cell depolarized at a similar rate. This demonstrates that the fragmentation we observe is not due to mitochondrial depolarization, nor does this artificially induced mitochondrial fragmentation cause subsequent loss of membrane potential.

**Figure 4.**
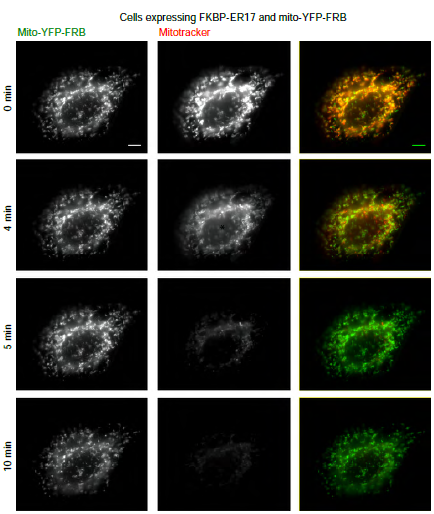
Addition of CCCP causes depolarization of mitochondrial fragments. Single-plane frames from time-lapse images of cells shown in Fig1D, E following addition of 100 nM CCCP. Scale bar = 10 μm.

To confirm that the mitochondrial fragments generated by inducible tethering via FKBP-ER17 were indeed individual elements and not contiguous with the rest of the mitochondrial network, we used fluorescence recovery after photobleaching (FRAP). Regions were bleached and afterwards imaged to detect signs of recovery in cells co-transfected with mito-YFP-FRB, FKBP-ER17 and DsRed-Mito plasmids and incubated with rapamycin for 1 hr prior to imaging (Fig. 5). To label the mitochondria independently of the mito-YFP-FRB fusion, we used DsRed-Mito, which is formed of the mitochondrial targeting sequence from subunit VIII of human cytochrome c oxidase fused to the red fluorescent protein DsRed. We bleached DsRed-mito selecting regions that appeared to be isolated (which we expected not to show signal recovery) in addition to connected regions (to confirm that recovery could occur). This approach enabled us to photobleach the DsRed-mito fusion without affecting the mito-YFP-FRB thereby allowing us to track the photobleached mitochondrial fragments using the YFP channel.

**Figure 5.**
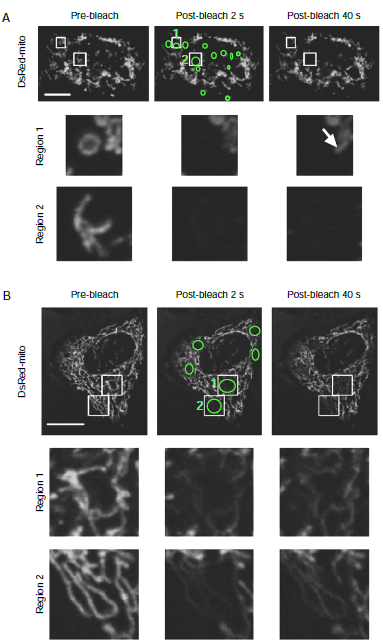
FRAP analysis of mitochondrial luminal continuity. DsRed-mito fluorescence in HeLa cells additionally co-transfected with mito-YFP-FRB (not shown) and FKBP-ER17 incubated with rapamycin for 1 hr. (A) Cell showing mitochondrial fragmentation phenotype. Cell was exposed to 5 pre-bleach frames and 6 bleach frames. In region 1 and region 2 isolated fragments do not show recovery. In region 1 a continuous area to the right is bleached and then recovers (arrow), while the curved mitochondrion in the middle does not. (B) Cell with continuous mitochondrial network, where regions show recovery 40 s post-bleaching. Cell exposed to 3 pre-bleach and 8 bleach frames. Frame rate 1 frame per 2 seconds. Scale bars = 10 μm.

We tested 96 regions over 20 cells; 60 regions that appeared to be isolated and 33 regions that appeared to be connected. 93% of connected regions showed recovery of fluorescent signal within 35s post bleach, while 98% of isolated regions did not.

Fig. 5A shows a cell with representative isolated regions, while Fig. 5B shows a cell with a fully-intact mitochondrial network. Despite the connected regions being bleached for a slightly longer time period in this instance (8 seconds compared to 6 for the isolated regions) recovery was not seen after 35 seconds, at which time point recovery was seen in the connected regions, and no recovery was seen even almost 2 minutes after bleaching (Movies 4 and 5). Full recovery of fluorescence was not seen within the time-scale of these experiments. This is consistent with previous work using FRAP on mitochondria (Collins and Bootman, 2003). These results demonstrate that the mitochondrial fragments generated by inducible tethering of the ER to the mitochondria are distinct entities isolated from the remainder of the mitochondrial network.

We then sought to determine whether this fission event was driven by close apposition of the two membranes. ER exit sites are continuous with the ER membrane but several of the components of the COPII budding machinery are localized directly adjacent to the ER membrane at these sites, rather than being contiguous with it. An example of this is Trk-fused gene (TFG) which associates tightly with the COPII budding machinery that drives secretory cargo exit from the ER and has been proposed to facilitate COPII assembly at ER exit sites (ERES) (Witte et al., 2011). ERES are spaced throughout the ER network and therefore could explain the relatively regular spacing of sites of fission seen in Figures 2 and 3. We sought to define whether we could trigger mitochondrial fission using an FKBP-TFG fusion.

When expressed in cells, FKBP-TFG forms discrete puncta similar to those seen with GFP-TFG ((Witte et al., 2011) and our unpublished data). These puncta co-localize with the ERES protein Sec31A (Fig. 6). Time-lapse imaging following addition of rapamycin revealed clustering of the mito-YFP-FRB (Fig. 7, Movie 6) when co-expressed with FKBP-TFG. However, we did not observe mitochondrial fragmentation, even 55 minutes after addition of rapamycin. As seen for FKBP-ER17, mitochondria did not depolarize until addition of CCCP and this then resulted in fading of the MitoTracker signal. We conclude that inducible tethering of ERES is not sufficient for mitochondrial fragmentation.

**Figure 6.**
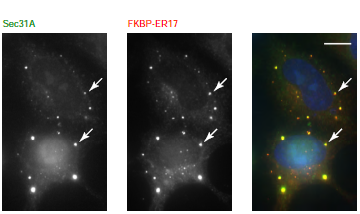
Localisation of FKBP-TFG. Immunofluorescence labelling of methanol fixed HeLa cells expressing FKBP-TFG. A dilute concentration of anti-TFG antibody was used to compensate for the over-expression of the FKBP-TFG protein. Arrows show puncta. Scale bar = 10 μm.

**Figure 7.**
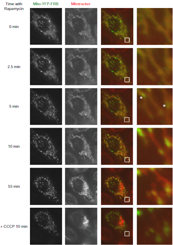
Inducible tethering of TFG to mitochondria. Single-plane frames from time-lapse imaging of HeLa cells co-transfected with FKBP-TFG and mito-YFP-FRB following addition of 200 nM rapamycin then 100 nM CCCP. Asterisks indicate clustering of mito-YFP-FRB. Scale bar = 10 μm.

## Discussion

Membrane-membrane interactions cross-link many organelles in eukaryotic cells, with interactions with the ER (as the largest organelle) being particularly frequent. Much recent interest has focused in the functional role of ER contacts including ER-mitochondrial calcium transfer (Csordas et al., 2010) and mitophagy (Bockler and Westermann, 2014) as well as their involvement in mitochondrial division (Friedman et al., 2011; Murley et al., 2013). Here we have demonstrated that inducible tethering of the ER membrane to mitochondria is sufficient to cause rapid mitochondrial fragmentation. We observe a two-step process here. Both FKBP-TFG and FKBP-FP17 induce a clustering of mito-YFP-FRB on the mitochondrial membrane but only FKBP-FP17 causes subsequent fragmentation of mitochondria. This kinetic delay could be due to assembly of the fission machinery or reorganization of the ER-mitochondria membrane interface to facilitate fission. Our interpretation of the fact that FKBP-TFG does not induce fission (even after several hours in the presence of rapamycin) is that its localization is simply not close enough to the ER membrane to direct close enough proximity of the two membranes to trigger the fission event or alternately that the presence of ERES is somehow incompatible with the processes required to facilitate fission.

We do not observe mitochondrial fission when using the FKBP-TFG fusion. Neither do we observe mitochondrial fission when artificially targeting the AP2 clathrin adaptor (data not shown and see (Robinson et al., 2010)). We tested the potential of an FKBP-TFG fusion to induce mitochondrial fragmentation when targeted to the outer mitochondrial membrane for two reasons. Both ER-mitochondrial contact sites have been implicated in the initiation of autophagy (Hamasaki et al., 2013) and ER exit sites have been shown to be important sites of autophagy initiation (Tan et al., 2013). We therefore postulated that ERES could act as a hub for these two processes. While we do not see mitochondrial fission with the FKBP-TFG fusion, it is important to note that on addition of rapamycin, FKBP-TFG does induce clustering of mito-YFP-FRB showing that this alone is not sufficient to drive mitochondrial fission. We also cannot rule out that the targeting and or function of FKBP-TFG is compromised compared to that of endogenous TFG. As such FKBP-TFG acts as a negative control here and we do not draw any firm conclusions with regard to the role that TFG or ERES might play in ER-mitochondrial contact.

One caveat that should be borne in mind when using the inducible systems currently available is the potential effects resulting from adding rapamycin to cells, as rapamycin is a potent inhibitor of the mTOR signalling pathway, to cells. mTOR co-ordinates protein turn-over and autophagy in response to nutrient availability (Raught et al., 2001). In these experiments, imaging of cells expressing FKBP-TFG and non-expressing cells demonstrates that the mitochondrial fragmentation we see with FKBP-ER17 is not due to the effects of rapamycin.

Since the ER membrane has previously been shown to be intimately involved in establishing the site of mitochondrial division in yeast and mammals (Friedman et al., 2011), we speculate that inducible tethering of the ER membrane may trigger fission through a similar mechanism. If simply bringing the ER and mitochondria into close proximity is sufficient to divide the mitochondrial network, one important question that remains is how are normal ER-mitochondrial contract sites regulated to prevent constitutive and uncontrolled mitochondrial division? In yeast, the cortical ER acts to tether mitochondria for segregation in the nascent bud (Swayne et al., 2011). How is this achieved without triggering mitochondrial fragmentation? Similarly, it is not clear why the number of sites of mitochondrial fission is limited following artificial tethering of ER and mitochondrial membranes. This could be due to the physical nature of mitochondria e.g. a finite size of the smallest fragment that we can generate owing to geometric constraints or organization of nucleoids (DNA within the mitochondria) that appear to localize adjacent to sites of fission (Ban-Ishihara et al., 2013; Murley et al., 2013). It could also be due to a limiting quantity of the fission machinery, or due to a defined number of pre-existing sites already primed for fission awaiting apposition of ER and mitochondrial membranes. Clearly there are many open questions here and we hope that a synthetic system to trigger time-resolved mitochondrial fission will be of value to study the machinery, regulation and role of ER-associated mitochondrial fission.

## Materials and Methods

### Plasmid Construction

pLVX.FKBP was constructed by using PCR to amplify the FKBP domain from the γ-FKBP plasmid (a kind gift from M. Robinson, Cambridge (Robinson et al., 2010)) and inserted into pLVX.puro (Clontech, California) using XhoI/EcoRI sites. This XhoI site was destroyed in the cloning and a novel XhoI site introduced downstream of the FKBP domain. Subsequently ER17 or TFG were amplified by PCR from GFP-FP-17 (the kind gift of N. Borgese, Milan) or GFP-TFG (the kind gift of A. Audhya, Wisconsin) and inserted into pLVX.FKBP using XhoI/XbaI or XhoI/BamHI respectively. Sequences amplified by PCR were subsequently confirmed by sequencing the resultant plasmid.

pMito-YFP-FRB and γ-FKBP construct were a kind gift of Scottie Robinson (Robinson et al., 2010) and pDsRed-Mito was obtained from Clontech.

### Cell Culture

HeLa cells were cultured in DMEM medium (Sigma-Aldrich, Poole, UK). Plasmids were transfected using Lipofectamine^®^ 2000 (Life Technologies, Paisley, UK) as per manufacturer’s instructions.

### Imaging

Glass-bottomed dishes (MatTek, Ashland, MA, USA) were used for live cell imaging. Where used, MitoTracker^®^ Red CMXRos (Life Technologies, Paisley, UK) was added to media at 1:2000 dilution for 3-5 min immediately prior to imaging, and then rinsed to remove excess dye. Cells were then placed in imaging media (Life Technologies, Paisley, UK). Rapamycin (Sigma-Aldrich, Poole, UK) was added as a 0.025 mg.ml^-1^ stock in DMSO to a final concentration of 200 nM. CCCP was added as a 1000x stock to a final concentration of 100 nM.

Live epifluorescence imaging was performed with an Olympus IX-81 inverted microscope fitted with a 37°C heated Perspex box using a 63× oil objective and an Orca-R2 CCD camera. Z-stacks are 9 × 1 micron slices; all time-lapse imaging is of a single focal plane. For immunofluorescence, cells were methanol-fixed prior to antibody staining. Mouse anti-FKBP12 and mouse anti-Sec31A were obtained from BD Transduction Laboratories, Oxford, UK; rabbit anti-TFG is from Imgenex, San Diego, USA. Fixed cells were imaged with an Olympus IX-71 inverted microscope. Volocity software (Perkin Elmer, version 5.4.2) was used for image acquisition and Image J (Fuji version 1.48q (Schindelin et al., 2012)) for image processing.

### Fluorescence Recovery after Photobleaching

Cells were live imaged in a 37°C heated Perspex box (Life Imaging Services, Reinach, CH) on a Leica SP5 confocal imaging system (Leica Microsystems, Milton Keynes, UK) with a 63x 1.4 numerical aperture blue light-corrected lens. EYFP was imaged using the 514 nm laser line and DsRed using the 594 nm laser line. A pinhole size of 1.8 Airy disk units was used to take images using a Leica DMI 6000 inverted microscope. Cells were exposed to 3-6 pre-bleach frames, 4-10 bleach frames with 594 nm laser at high power and at least 35 post-bleach frames at low laser power all at 1 frame per 2 seconds. Data was processed using Leica LAS AF Lite (Wetzlar, Germany) and Image J (Fuji version 1.48q (Schindelin et al., 2012)).

## Acknowledgments

We would like to thank Margaret Robinson, Nica Borgese, Anjon Audhya and Jon Lane for reagents. We are grateful to Jon Lane and Tom MacVicar for useful discussion and advice, also to Anna Brown for preliminary experimental work and to Niamh Nugent for assisting with aspects of experimental work. Ours thanks go to the MRC, Wolfson Foundation, and University of Bristol for supporting the Bristol Bioimaging facility used for the FRAP experiments.

## Contributions

V.J.M performed experimental work, analyzed the data and wrote the manuscript; D.J.S. conceived and directed the project, assisted with experimental work, and helped draft the manuscript.

## References

Alexander, C., M. Votruba, U.E. Pesch, D.L. Thiselton, S. Mayer, A. Moore, M. Rodriguez, U. Kellner, B. Leo-Kottler, G. Auburger, S.S. Bhattacharya, and B. Wissinger. 2000. OPA1, encoding a dynamin-related GTPase, is mutated in autosomal dominant optic atrophy linked to chromosome 3q28. Nat. Genet. 26:211–215.

Ban-Ishihara, R., T. Ishihara, N. Sasaki, K. Mihara, and N. Ishihara. 2013. Dynamics of nucleoid structure regulated by mitochondrial fission contributes to cristae reformation and release of cytochrome c. Proc. Natl. Acad. Sci. USA. 110:11863–11868.

Benard, G., N. Bellance, D. James, P. Parrone, H. Fernandez, T. Letellier, and R. Rossignol. 2007. Mitochondrial bioenergetics and structural network organization. J. Cell Sci. 120:838–848.

Bereiter-Hahn, J., and M. Voth. 1994. Dynamics of mitochondria in living cells: shape changes, dislocations, fusion, and fission of mitochondria. Microscopy research and technique. 27:198–219.

Bockler, S., and B. Westermann. 2014. Mitochondrial ER Contacts Are Crucial for Mitophagy in Yeast. Dev. Cell. 28:450–458.

Brandizzi, F., and C. Barlowe. 2013. Organization of the ER-Golgi interface for membrane traffic control. Nature reviews. Molecular cell biology. 14:382–392.

Bulbarelli, A., T. Sprocati, M. Barberi, E. Pedrazzini, and N. Borgese. 2002. Trafficking of tail-anchored proteins: transport from the endoplasmic reticulum to the plasma membrane and sorting between surface domains in polarised epithelial cells. J. Cell Sci. 115:1689–1702.

Collins, T.J., and M.D. Bootman. 2003. Mitochondria are morphologically heterogeneous within cells. The Journal of experimental biology. 206:1993–2000.

Csordas, G., P. Varnai, T. Golenar, S. Roy, G. Purkins, T.G. Schneider, T. Balla, and G. Hajnoczky. 2010. Imaging interorganelle contacts and local calcium dynamics at the ER-mitochondrial interface. Molecular cell. 39:121–132.

Delettre, C., G. Lenaers, J.M. Griffoin, N. Gigarel, C. Lorenzo, P. Belenguer, L. Pelloquin, J. Grosgeorge, C. Turc-Carel, E. Perret, C. Astarie-Dequeker, L. Lasquellec, B. Arnaud, B. Ducommun, J. Kaplan, and C.P. Hamel. 2000. Nuclear gene OPA1, encoding a mitochondrial dynamin-related protein, is mutated in dominant optic atrophy. Nat. Genet. 26:207–210.

Deng, H., M.W. Dodson, H. Huang, and M. Guo. 2008. The Parkinson’s disease genes pink1 and parkin promote mitochondrial fission and/or inhibit fusion in Drosophila. Proc. Natl. Acad. Sci. USA. 105:14503–14508.

Friedman, J.R., L.L. Lackner, M. West, J.R. Dibenedetto, J. Nunnari, and G.K. Voeltz. 2011. ER Tubules Mark Sites of Mitochondrial Division. Science. 334:358–362.

Ge, L., D. Melville, M. Zhang, and R. Schekman. 2013. The ER-Golgi intermediate compartment is a key membrane source for the LC3 lipidation step of autophagosome biogenesis. Elife. 2:e00947.

Hackenbrock, C.R. 1966. Ultrastructural bases for metabolically linked mechanical activity in mitochondria. I. Reversible ultrastructural changes with change in metabolic steady state in isolated liver mitochondria. The Journal of cell biology. 30:269–297.

Hackenbrock, C.R. 1968. Ultrastructural bases for metabolically linked mechanical activity in mitochondria. II. Electron transport-linked ultrastructural transformations in mitochondria. The Journal of cell biology. 37:345–369.

Hamasaki, M., N. Furuta, A. Matsuda, A. Nezu, A. Yamamoto, N. Fujita, H. Oomori, T. Noda, T. Haraguchi, Y. Hiraoka, A. Amano, and T. Yoshimori. 2013. Autophagosomes form at ER-mitochondria contact sites. Nature. 495:389–393.

Hermann, G.J., and J.M. Shaw. 1998. Mitochondrial dynamics in yeast. Annual review of cell and developmental biology. 14:265–303.

Inoue, T., W.D. Heo, J.S. Grimley, T.J. Wandless, and T. Meyer. 2005. An inducible translocation strategy to rapidly activate and inhibit small GTPase signaling pathways. Nature methods. 2:415–418.

Kornmann, B., E. Currie, S.R. Collins, M. Schuldiner, J. Nunnari, J.S. Weissman, and P. Walter. 2009. An ER-mitochondria tethering complex revealed by a synthetic biology screen. Science. 325:477–481.

Korobova, F., T.J. Gauvin, and H.N. Higgs. 2014. A Role for Myosin II in Mammalian Mitochondrial Fission. Current biology : CB. 24:409–414.

Korobova, F., V. Ramabhadran, and H.N. Higgs. 2013. An actin-dependent step in mitochondrial fission mediated by the ER-associated formin INF2. Science. 339:464–467.

Lackner, L.L., and J.M. Nunnari. 2009. The molecular mechanism and cellular functions of mitochondrial division. Biochimica et biophysica acta. 1792:1138–1144.

Murley, A., L.L. Lackner, C. Osman, M. West, G.K. Voeltz, P. Walter, and J. Nunnari. 2013. ER-associated mitochondrial division links the distribution of mitochondria and mitochondrial DNA in yeast. Elife. 2:e00422.

Nguyen, T.T., A. Lewandowska, J.Y. Choi, D.F. Markgraf, M. Junker, M. Bilgin, C.S. Ejsing, D.R. Voelker, T.A. Rapoport, and J.M. Shaw. 2012. Gem1 and ERMES do not directly affect phosphatidylserine transport from ER to mitochondria or mitochondrial inheritance. Traffic. 13:880–890.

Nunnari, J., and A. Suomalainen. 2012. Mitochondria: in sickness and in health. Cell. 148:1145–1159.

Qian, W., S. Choi, G.A. Gibson, S.C. Watkins, C.J. Bakkenist, and B. Van Houten. 2012. Mitochondrial hyperfusion induced by loss of the fission protein Drp1 causes ATM-dependent G2/M arrest and aneuploidy through DNA replication stress. J. Cell Sci. 125:5745–5757.

Raught, B., A.C. Gingras, and N. Sonenberg. 2001. The target of rapamycin (TOR) proteins. Proceedings of the National Academy of Sciences of the United States of America. 98:7037–7044.

Robinson, M.S., D.A. Sahlender, and S.D. Foster. 2010. Rapid inactivation of proteins by rapamycin-induced rerouting to mitochondria. Dev. Cell. 18:324–331.

Ronchi, P., S. Colombo, M. Francolini, and N. Borgese. 2008. Transmembrane domain-dependent partitioning of membrane proteins within the endoplasmic reticulum. J. Cell Biol. 181:105–118.

Schindelin, J., I. Arganda-Carreras, E. Frise, V. Kaynig, M. Longair, T. Pietzsch, S. Preibisch, C. Rueden, S. Saalfeld, B. Schmid, J.Y. Tinevez, D.J. White, V. Hartenstein, K. Eliceiri, P. Tomancak, and A. Cardona. 2012. Fiji: an open-source platform for biological-image analysis. Nature Methods. 9:676–682.

Swayne, T.C., C. Zhou, I.R. Boldogh, J.K. Charalel, J.R. McFaline-Figueroa, S. Thoms, C. Yang, G. Leung, J. McInnes, R. Erdmann, and L.A. Pon. 2011. Role for cER and Mmr1p in anchorage of mitochondria at sites of polarized surface growth in budding yeast. Current biology : CB. 21:1994–1999.

Tan, D., Y. Cai, J. Wang, J. Zhang, S. Menon, H.T. Chou, S. Ferro-Novick, K.M. Reinisch, and T. Walz. 2013. The EM structure of the TRAPPIII complex leads to the identification of a requirement for COPII vesicles on the macroautophagy pathway. Proc. Natl. Acad. Sci. USA. 110:19432–19437.

Waterham, H.R., J. Koster, C.W. van Roermund, P.A. Mooyer, R.J. Wanders, and J.V. Leonard. 2007. A lethal defect of mitochondrial and peroxisomal fission. The New England journal of medicine. 356:1736–1741.

Witte, K., A.L. Schuh, J. Hegermann, A. Sarkeshik, J.R. Mayers, K. Schwarze, J.R. Yates Iii, S. Eimer, and A. Audhya. 2011. TFG-1 function in protein secretion and oncogenesis. Nat. Cell Biol. 13:550–558.

Yang, Y., Y. Ouyang, L. Yang, M.F. Beal, A. McQuibban, H. Vogel, and B. Lu. 2008. Pink1 regulates mitochondrial dynamics through interaction with the fission/fusion machinery. Proceedings of the National Academy of Sciences of the United States of America. 105:7070–7075.

Youle, R.J., and A.M. van der Bliek. 2012. Mitochondrial fission, fusion, and stress. Science. 337:1062–1065.

Zuchner, S., I.V. Mersiyanova, M. Muglia, N. Bissar-Tadmouri, J. Rochelle, E.L. Dadali, M. Zappia, E. Nelis, A. Patitucci, J. Senderek, Y. Parman, O. Evgrafov, P.D. Jonghe, Y. Takahashi, S. Tsuji, M.A. Pericak-Vance, A. Quattrone, E. Battaloglu, A.V. Polyakov, V. Timmerman, J.M. Schroder, and J.M. Vance. 2004. Mutations in the mitochondrial GTPase mitofusin 2 cause Charcot-Marie-Tooth neuropathy type 2A. Nat. Genet. 36:449–451.

